# Comprehensive sequencing of the lung neuroimmune landscape in response to asthmatic induction

**DOI:** 10.1101/2024.10.04.616730

**Authors:** Hayden McSwiggin, Rui Wang, Rubens Daniel Miserani Magalhães, Fengli Zhu, Taylor A. Doherty, Wei Yan, Nicholas Jendzjowsky

**Affiliations:** The Lundquist Institute for Biomedical Innovation at Harbor-UCLA, Torrance, CA 90502, USA; Division of Allergy and Immunology, Department of Medicine, University of California San Diego, Veterans Affairs San Diego Healthcare System, La Jolla, CA 92093, USA; Division of Endocrinology, Department of Medicine, Harbor-UCLA Medical Center Torrance, CA 90502, USA; Division of Respiratory and Critical Care Medicine and Physiology, Department of Medicine, Harbor-UCLA Medical Center, Torrance, CA 90502, USA

**Keywords:** Vagus, asthma, *Alternaria alternata*, RNA-seq, single-nucleus RNA-seq, spatial RNA-seq, transcriptomics

## Abstract

Evidence demonstrates that sensory neurons respond to pathogenic/allergic infiltration and mediate immune responses, forming an integral part of host defense that become hypersensitized during allergy. Our objective was to investigate how asthmatic induction alters the pulmonary neuroimmune transcriptome. We hypothesized that asthmatic induction would upregulate genes in the vagal ganglia (nodose/jugular ganglia), which would be associated with asthmatic immunity, and that these would be clustered, primarily in nodose neurons. Furthermore, lungs would increase transcripts associated with nerve activation, and these would be centered in neural and neuroendocrine-like cells. Bulk RNA-seq revealed that genes related to allergen sensing were increased in asthmatic ganglia nodose/jugular ganglia compared to control ganglia. These genes were associated with nodose clusters as shown by single-nucleus RNA sequencing, and a distinct caudal-to-rostral spatial arrangement was presented as delineated by spatial transcriptomics. The distinct clusters closely match previous identification of nodose neuron clusters. Correspondingly, the lung transcriptome was altered with asthmatic induction such that transcripts associated with neural excitation were upregulated. The spatial distribution of these transcripts was revealed by spatial transcriptomics to illustrate that these were expressed in neuroendocrine-like cells/club cells, and neurons. These results show that the neuroimmune transcriptome is altered in response to asthmatic induction in a cell cluster and spatially distinct manner.

**Significance statement:** This study provides a comprehensive transcriptomic map of the neuroimmune alterations in response to asthmatic induction.

## Introduction

Asthma is characterized by variable airflow obstruction where exaggerated airway narrowing occurs due to aberrant inflammation in response to innocuous substances(1). Thus, inflammation is central to the pathogenesis of asthma(2). The surveillance of the lung by vagal afferent nerves is tasked to protect the lung from foreign substances in the form of chemosensitive and mechanosensitive reflexes(3–5). Vagal sensory neuron hypersensitization is known to induce quick airway reflexes such as cough and bronchoconstriction(3, 4) in response to their stimulation, which is both dependent on and independent of immune cell-mediated stimulation(6). Vagal sensory afferent nerves also play an important role in driving asthmatic airway hyperresponsiveness and inflammation(4). Specifically, sensory neurons are sensitized by cytokines(7, 8), pathogens(9, 10), and allergens(11, 12) which release neuropeptides that activate T cells to induce inflammation(8, 13), B cells to release immunoglobulins(14), mast cells to degranulate(15, 16) or traffic eosinophils(8, 13). Therefore, the interplay between sensory neurons, stromal cells, and immune cells dictates asthmatic pathogenesis.

Neuron sub-types can be classified by genomic features that directly reflect both ontogeny and function, with a taxonomy containing four principal types: cold, mechano-heat, A-low threshold mechanoreceptors, and mechano-heat-itch and C-low threshold mechanoreceptors where each type branches into several neuron sub-types with unique response properties(17). This heterogeneity among neuron types is the reason for the cellular basis for discrimination among somatic sensory modalities and forms the basis for the ability to perceive, explore, and interpret the internal organ function(18, 19). Recent transcriptomic analyses have allowed the identification of distinct neural subsets involved in interoception, chemosensation, mechanosensation, and a host of homeostatic processes(18, 19) including single-cell RNA-seq analysis which demonstrates molecularly unique subtypes of vagal sensory neurons(3, 18). Furthermore, molecularly distinct subsets of vagal sensory neurons have specific nerve terminals depending on their sensory modalities and functions(19, 20) and their functional properties are largely determined by their molecular identity(3, 20). This provides important information to predict possible phenotypic changes to vagal immune sensing ganglia in response to disease. To date, however, few data are available to elucidate how vagal ganglia and respective target organs are transcriptionally altered in response to disease states. This information would be valuable for deterministic investigation of the mechanistic underpinnings of asthmatic inflammation.

In this study, we were motivated to investigate the simultaneous phenotypic changes induced by asthmatic sensitization to the vagal ganglia and lungs. Driven by previous investigations that provided distinct delineation of vagal sensory neuron subsets, we wanted to provide a complete transcriptomic profile of vagal sensory neurons in response to asthmatic induction along with lung-associated transcriptomic changes. We show that, globally, the lungs appear to upregulate neural transcription factors whereas the vagal sensory afferent upregulates immune transcription factors. We used ‘bulk’ RNA sequencing of lungs and vagal ganglia, single-nucleus RNA sequencing of paired vagal ganglia, and spatial transcriptomic profiling of asthmatic lungs and vagal ganglia to showcase how asthmatic induction alters the neuro-immune system of the lungs.

## Results

### Confirmation of asthmatic induction

Mice subject to *A. alternata* sensitization displayed enhanced goblet cell metaplasia and heightened IgE concentrations in serum (**Figure 1A-D**) compared to PBS controls, demonstrating the induction of an allergic asthmatic phenotype consistent with our previous results(14, 21).

**Figure 1.**
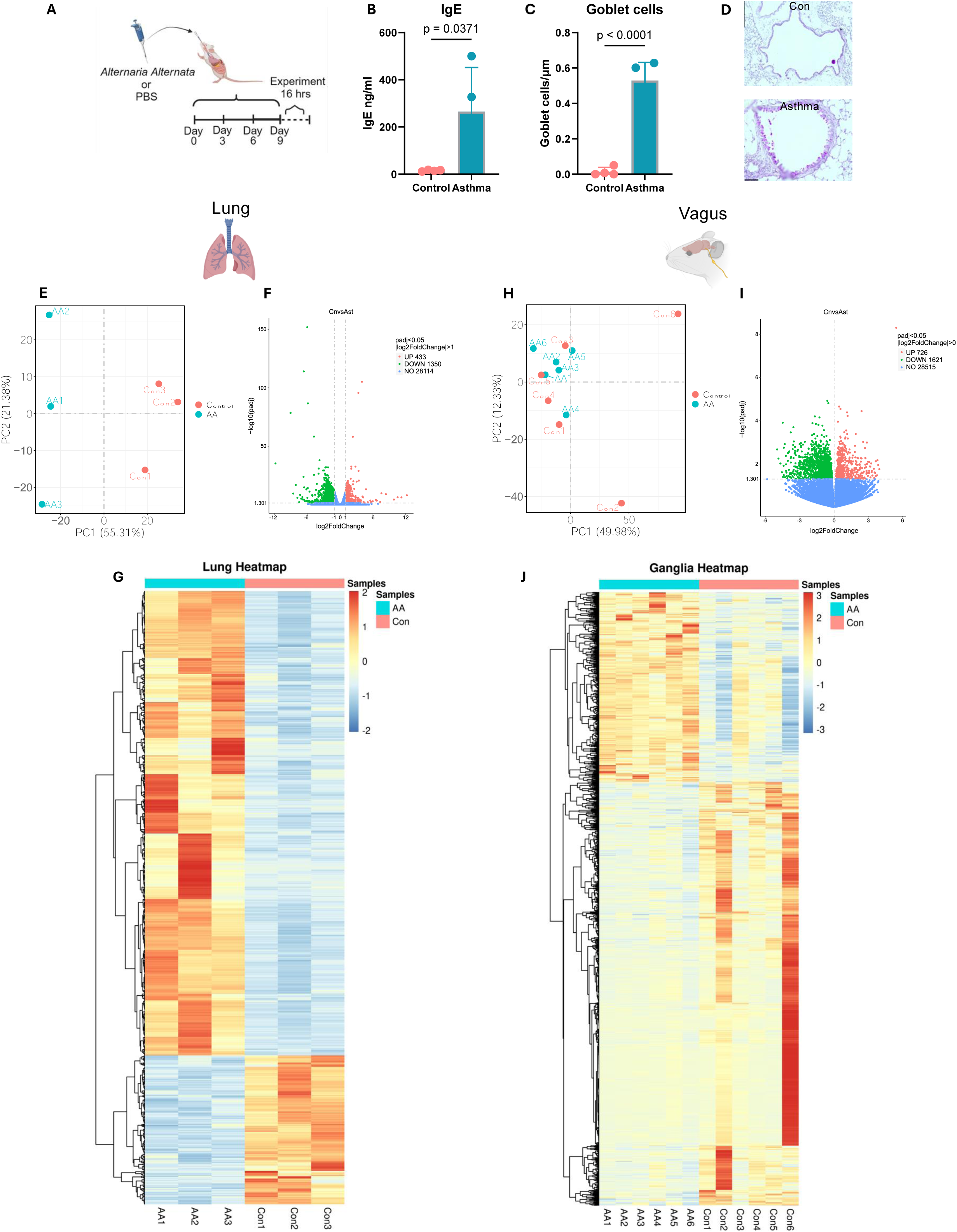
*Alternaria alternata* alters the neuroimmune transcriptome. **A)** Model of *A. alternata* asthmatic induction(14, 48). Image created with Biorender. **B)** Serum levels of IgE were elevated after induction of asthma with *A. alternata* (Two-sided t-test). **C)** Goblet cell metaplasia was induced with *A. alternata* (Two-sided t-test). N=4 per group. **D)** Representative images, 20x, scale bar= 20µm. **E)** PCA analysis shows distinct clustering patterns based on allergic induction in lungs. **F)** Volcano plot displaying differentially expressed genes from lungs (x-axis represents log2FC in gene expression, y-axis represents –log10 adjusted p-value). **G)** Heatmap of differentially expressed lung transcripts between control and *A. alternata* groups. **H)** PCA analysis showing difference between gender and allergic induction in vagus. **I)** Volcano plot displaying differentially expressed vagus genes (x-axis represents log2FC in gene expression, y-axis represents –log10 adjusted p-value). **J)** Heatmap of differentially expressed vagus transcripts between control and *A. alternata* groups.

### Lung transcriptome

Bulk sequencing of whole lungs showed an average of 54,922,129 reads of 29,897 genes across samples. Principle component analysis revealed that 56% and 21% were sufficient to describe the differences between samples (**Figure 1E**). We then performed a two-group comparison of each ganglia group to the other two groups and identified the genes that were enriched, or depleted, DESeq2, p≤ 0.05. Applying these criteria, we identified 1783 unique genes that were differentially regulated in *A. alternata* mice compared to PBS which included a downregulation of 433 genes and upregulation of 1350 genes compared (**Figure 1F**). Gene expression differences are displayed as heatmaps in **Figure 1G**. Of note, the *Mrgprg* was significantly increased (Log2FC=6.999, p_adj_=3.81*10^−8^). A full list of these genes with their transcript LogF2C numbers are shown in **Supplementary Data S1**.

Next, we used Gene Ontology (GO), and KEGG orthology analysis to identify likely gene sets which were differentially expressed in *A alternata* lungs compared to PBS control lungs. GO analysis showed a differential upregulation of eosinophil migration (GO:0072677, p_adj_=0.000337), eosinophil chemotaxis (GO:0048245, p_adj_=0.000458), and mucosal immune response (GO:0002385, p_adj_=0.00152) demonstrated a robust asthmatic phenotype. Interestingly, upregulation of response to pain (GO:0048265, p_adj_=0.001067), behavioral response to pain (GO:0048266, p_adj_=0.009631) showed an augmented upregulation of sensory neuron genes (**Supplementary figure 1A**). KEGG orthology revealed a differential regulation of neuroactive ligand-receptor interaction (mmu04080, p_adj_=6.08×10^−7^) along with calcium signaling pathway (mmu04020, p_adj_=0.002555), asthma (mmu05310, p_adj_=0.000791) and cytokine-cytokine receptor interaction (mmu04060, p_adj_=0.0000359), **Supplementary figure 1B**).

### Vagal transcriptome

Bulk sequencing of whole vagal ganglia showed an average of 55,046,103 reads of 33,423 genes across all samples. The two principle components (sex, vs disease state) revealed that 50% and 12% were sufficient to describe the differences between samples (**Figure 1H**). We then performed a two-group comparison of each ganglia group to the other two groups and identified the genes that were enriched, or depleted. Transcripts were filtered by using the p-value, p ≤ 0.001. Applying these criteria, we identified 2347 unique differentially regulated genes in *A alternata* mice compared to PBS with an upregulation of 726 and a downregulation of 1621 genes compared to PBS mice (**Figure 1I**). Again, *Mrgprg* (Log2FC= 5.44, p_adj_=1.52*10^−4^) and other mas related GPCRs were upregulated in the vagal transcriptome Gene expression differences are displayed as heatmaps in **Figure 1J**.

Next, we used GO and KEGG orthology to identify likely gene sets that were differentially expressed in *A alternata* ganglia compared to PBS control ganglia. Specific GO analysis revealed that transcripts associated with immunogenic processes were upregulated in the vagal ganglia, such as cytokine production involved in immune response (GO:0002367, p_adj_=0.000015), and cytokine-mediated signaling pathway (GO: GO:0019221, p_adj_=0.0000013) (**Supplementary figure 1C**). KEGG orthology revealed a single significant upregulation of Neuroactive ligand receptor (mmu04080, p_adj_= 0.009449; **Supplementary figure 1D**). A full list of these genes with their transcript LogF2C numbers are shown in **Supplementary Data S2**.

## Single-nucleus RNA sequencing

A total of 2,640 cells from *A. alternata* and 5,030 cells from PBS-treated mice were initially analyzed. After quality control (QC) to remove low-quality cells and potential doublet cells, 1,627 from *A. alternata* and 3,416 cells from PBS-treated mice passed the QC requirements. After feature selection, removal of confounding sources of variation, and principal-component analysis, the cells were clustered by the Seurat Louvain-based algorithm. The first round of analysis at a resolution of 0.5 produced 13 clusters, and after merging highly similar groups, 11 clearly distinctive cell types remained (**Figure 2A, Supplementary figure 2**). All neuron clusters included cells from each biological replicate, demonstrating that the transcriptional profiles were robust and reproducible over the >5000 number of cells collected between 2 separate experiments and groups (**Figure 2B-D**).

**Figure 2.**
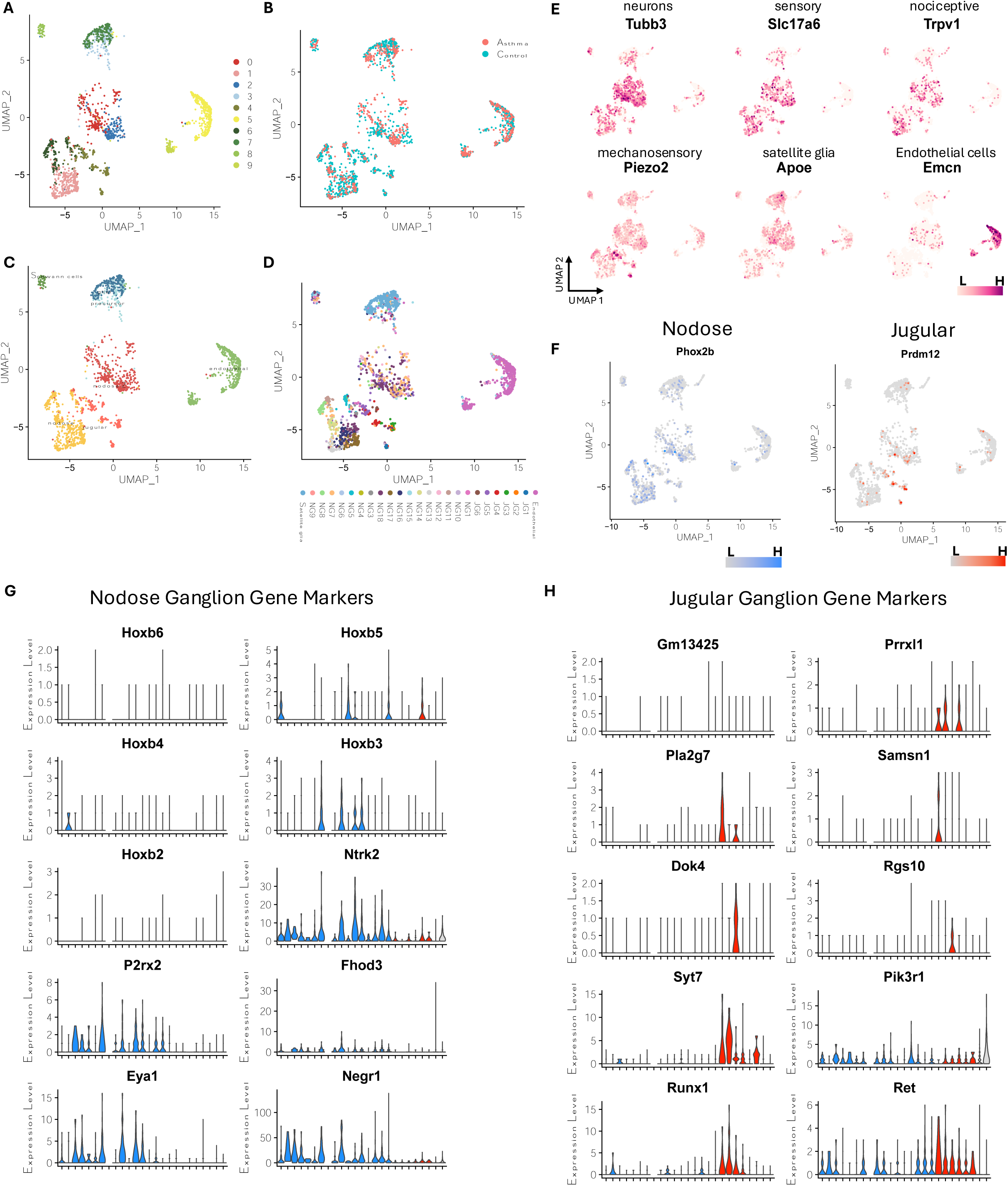
Single-nucleus RNA-seq analysis shows *Alternaria alternata* produces increased sensory transcripts in the vagal ganglia. **A)** Unbiased clustering reveals 11 distinct clusters in vagal ganglia in control and asthmatic ganglia. **B)** UMAP projection of vagal ganglia cells from asthmatic and control conditions **C)** General cellular annotation using canonical gene markers. **D**) Specific cellular annotation using RCTD method with Kupari et al. dataset as the reference. **E)** Canonical gene markers used to identify general cell types. **F)** Prdm12 and Phox2b delineate jugular and nodose cell subsets. **G**) Key nodose ganglion gene markers identified by Kupari et al. are prominent shown in nodose clusters. **H)** Key jugular ganglion gene markers identified by Kupari et al.(18) are prominent shown in jugular clusters.

Of the 11 clusters, 5 clusters expressed neuronal marker genes, including tubulin Beta 3 (Tubb3), synaptosome associated protein 25 (Snap25), ELAV-like binding protein 4 (*Elavl4*), and RNA binging fox-1 homolog 3 (*Rbfox3*). These clusters also expressed either the nodose marker *Phox2b* or the jugular marker *Prdm12* (**Figure 2E, F**) which are established neuronal transcripts(18, 22). Importantly, *Phox2b*^*+*^ and *Prdrm12*^*+*^ neurons were distinctly separated and contained unique gene sets (**Figure 2E-H**), closely corroborating previous findings despite the use of different methodologies (single-cell vs single-nucleus) (**Supplementary figure 3 & 4D**). For instance, cluster 4 which showed higher expression of *Prdm12* also showed higher expression of *Prrxl1, Samsn1*, and *Syt7*. Clusters 1 and 6, which are also *Phox2b*^*+*^, show expression of other nodose marker genes such as *Hoxb3, Ntrk2, P2rx2, Fhod3, Eya1*, and *Negr1*. Clusters 0 and 2 have many nodose neuron makers but also show specific expression of *Hoxb5*. Cluster 8 showed higher expression of genes marking Schwann cells such as *Pmp22, Sox10*, periaxin (*Prx*), and myelin protein zero (*Mpz*) myelin-associated protein (*Ncmap*). Cluster 7 showed higher expression of glial cell marker genes: *Aspa*, which plays a role in the metabolism of myelin, in addition to Lama2, Cdh19, and *Abca8a* (**Figure 2G, H**), indicating this may be satellite glial cells (SGCs). Cluster 3 may represent a precursor or transitional state leading to cluster 7 (SGCs) as they share many marker genes, but cluster 3 shows expression of marker genes that are typically associated with cell differentiation such as *Adgrb3* and *Lpar1*. Additionally, clusters 5 and 9 expressed endothelial cell(EC) marker genes: endomucin (*Emcn*), platelet and endothelial cell adhesion molecule 1 (*Pecam1*), and friend leukemia integration1 (*Fli1*). Cluster 5 expresses genes involved in angiogenesis and vascular integrity such as *Slco1a4, Abcg2, Lef1, Fli1*, and *Flt1* which suggest these may be vascular endothelial cells. Cluster 9 may represent a more specialized subset of metabolically active endothelial cells, potentially involved in lipid metabolism or related to vasculature of adipose tissues due to the higher expression of genes such as *Cd36, Fabp4*, and *Pparg*. To further annotated the cell types, we utilized a highly detailed single-cell RNA-seq dataset published by Kupari et. al.(18) which resulted in the annotation shown in **Figure 2D**.

A total of 222 genes identified as differentially expressed in the bulk RNA-seq data were also found to be differentially expressed between control and asthma conditions in the nodose and jugular of the single-nuclei RNA-seq data. Many of these genes have previously linked to inflammation and may play roles in the response to asthma. The key difference between clusters are shown in **Supplementary Data S3**). For example *Tgfbr2* and *Tgfbr3* were both down regulated in both dataset and were shown to be expressed in the jugular neurons in the single-nuclei dataset. Additionally, *B2m, Cd36, Epas1, Hspg2, Itga1*, and *Adgrf5* were downregulated in both dataset and specifically in jugular and nodose neurons, all of which are known to be involved in immune responses. Other genes previously shown to play roles in inflammation or asthma condition were upregulated in both dataset such as *Adnp, Arap2, Jund, Mxra7* and *Slc39a11*.

Using CellChat, we identified 1260 and 1416 interactions between control and asthma (**Figure 3A**) which are grouped into 56 pathways (**Figure 3B**). CellChat further revealed 156 additional interactions with higher association that were grouped into 12 different pathways (**Figure 3C, D**). Those pathways include those with key roles in tissue repair, fibrosis, and airway remodeling, such as BMPa, FGF, CDH, and COLLAGEN, which are crucial processes in chronic asthma(23–26). Other interactions were linked to NRG and SEMA3 which are involved in neuronal signaling, inflammation, and immune regulation, all of which are integral to asthma pathophysiology(27–30). TENASCIN, AGRN, and CADM contribute to tissue and synapse remodeling, likely affecting airway structure and neurogenic inflammation(31–34). Glutamate and DHEA are linked to neuronal signaling and immune modulation, influencing bronchoconstriction and inflammation in asthma(35–39). Lastly, ADGRG is involved in immune cell migration and activation, impacting the immune response during asthma hypersensitivity(40–42). Together, these pathways were shown to have more interactions in the asthma group, which contribute to the inflammation, immune response, and structural changes in asthma, particularly in response to allergens like *A. alternata*. Cell types with the most significant changes in sending or receiving signals after asthma induction were predominant in jugular and nodose cells (**Figure 3D**). One of the prominent CellChat pathways upregulated with asthma induction was neuregulin (NRG) (**Figure 3B, C**). This pathway is increased with nodose and jugular neurons being the primary senders to Schwann cells, with glial cells as influencers (**Figure 3E**). Differentially regulated ligands-receptors pairs between conditions, with jugular and nodose neurons as the source and other neurons are the target, are shown in **Supplementary Figure 5**.

**Figure 3.**
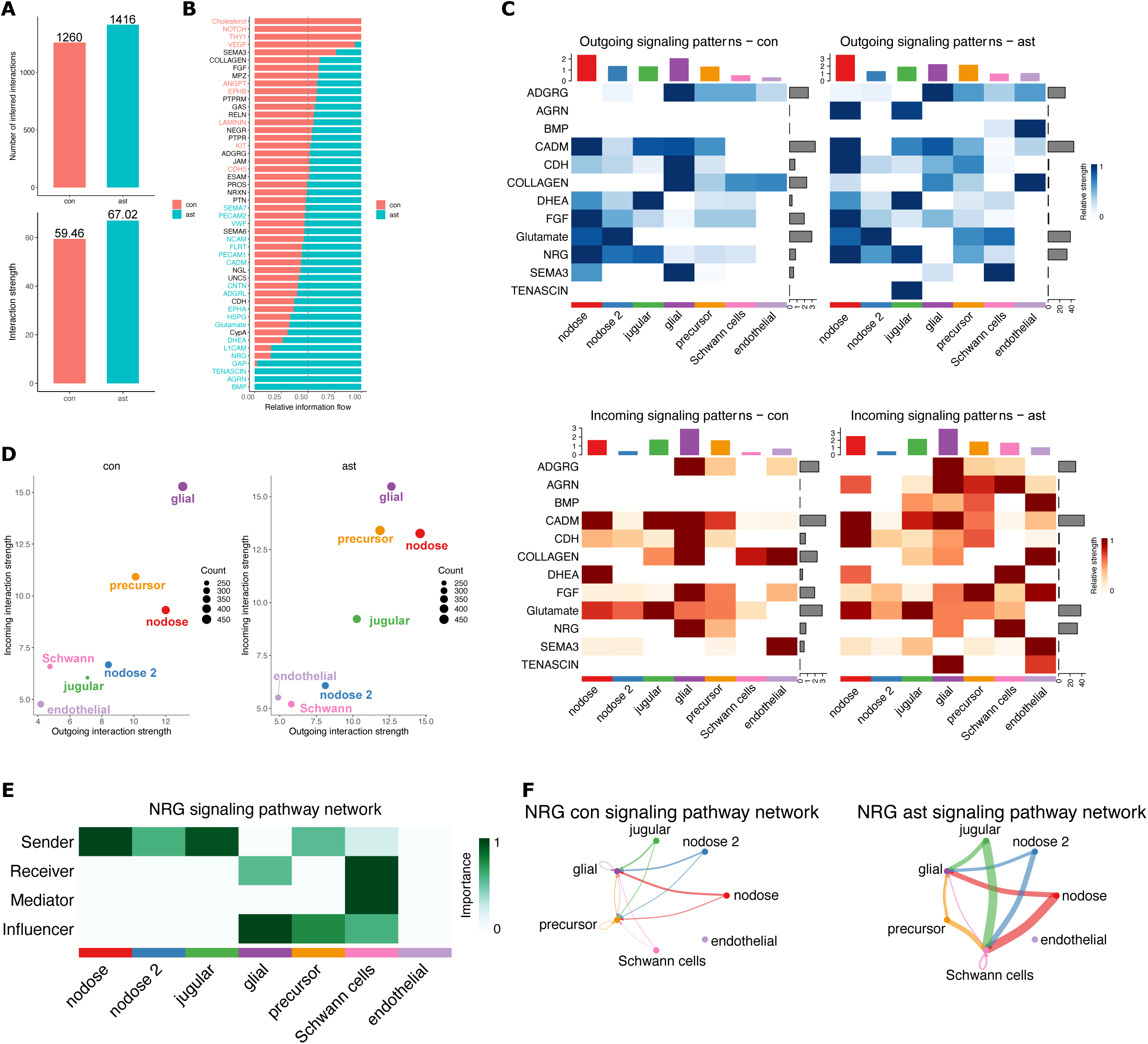
CellChat analysis. **a)** Number and strength of interactions identified by CellChat **b)** Information flow of each signaling pathway between *A. alternata* and control vagal neurons. **c)** Significant outgoing signals from neuron related pathways between *A. alternata* and control vagal neurons. **d)** Cell populations with significant changes in sending or receiving signals between *A. alternata* and control vagal neurons. **e)** NRG signaling pathway network showing cell populations in that are sending, receiving, mediating and influencing after *A. alternata* exposure (top) and the change in cell-to-cell communication with regards to NRG between *A. alternata* and control vagal neurons (bottom).

## Spatial sequencing

The selected 493 genes included 300 genes provided by the Xenium Mouse Tissue Atlassing panel and 92 custom genes selected from genes of interest from bulk sequencing and single-nucleus sequencing (**Supplementary Data S4**). Of the analyzed genes, we identified vagal cell clusters from those identified in single-nucleus sequencing (**Figure 4A**). The genes identified are consistent with upregulation in both snRNA-seq and bulk RNA-seq data where the agreement is (**Supplementary Figure 4C**). The clusters identified in single-nucleus sequencing were then mapped spatially to identify their tissue distribution which confirms the spatial distribution confirms the clustering of cell types as presented previously(3, 18) (**Figure 4B**). These clusters are reminiscent of all clusters identified from snRNA-seq (**Figure 4B**).

**Figure 4.**
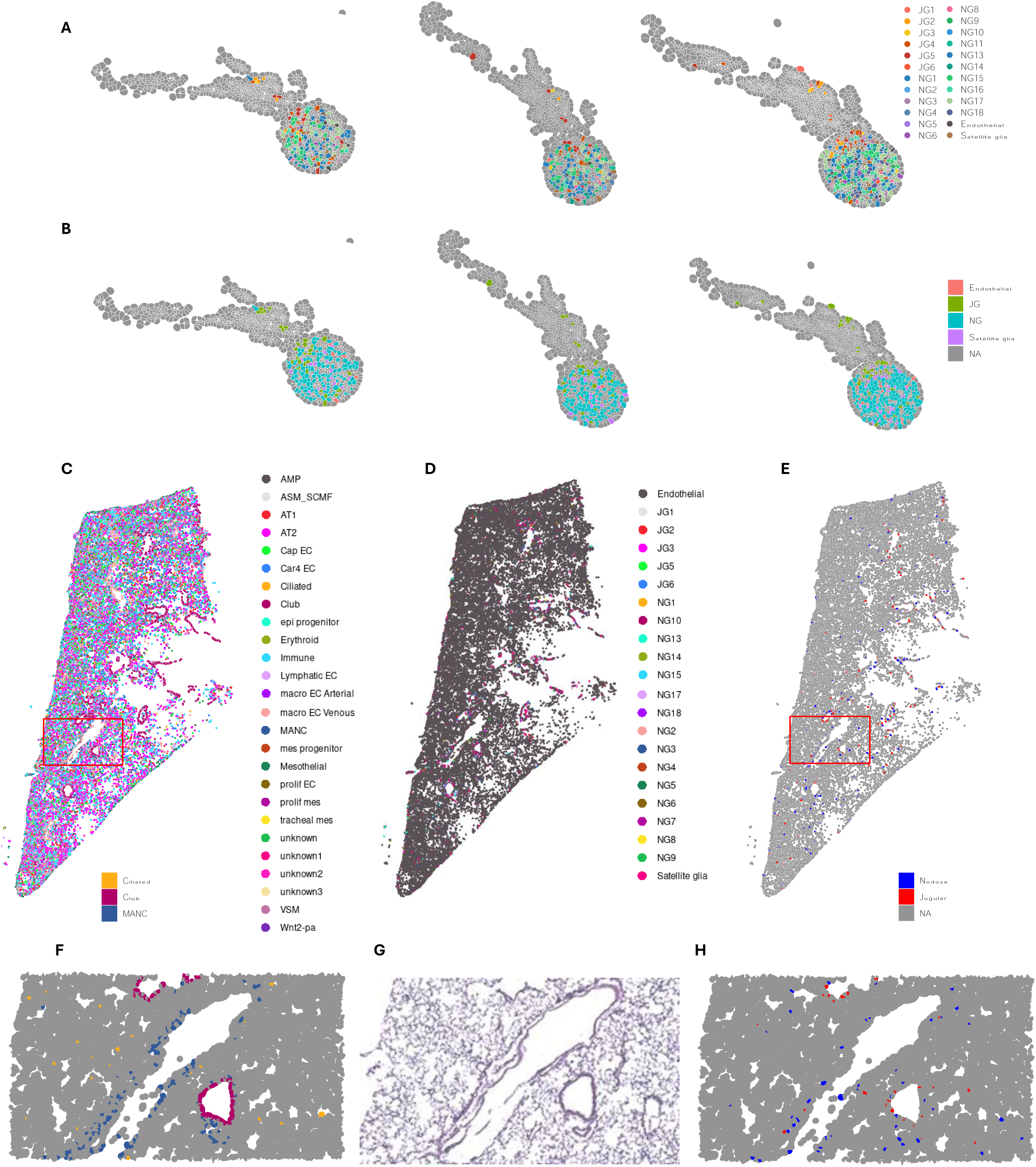
Spatial distribution of vagal transcriptome in the ganglia and lungs. **A)** Cell types identified by single nucleus sequencing, projected onto spatial map of vagus. **B**) Location of jugular and nodose cell types in vagus. **C**) Spatial lung data was deconvoluted using LungMAP single-cell dataset to identify all cell types and projected onto UMAP coordinates. **D**) Lund spatial dataset deconvoluted with the current vagal single-nucleus and Kupari dataset. **D)** Spatial distribution of jugular and nodose clusters. **F)** Enlarged area outlined in red from panel **C** showing spatial distribution of ciliated, club, and MANC cells. **G)** Enlarged H&E stain of lung section. **H)** Enlarged area outlined in red from panel **E** showing the spatial distribution of nodose and jugular cells.

We then used a published lung single-cell dataset to deconvolute the spatial lung dataset (**Figure 4C**). When we superimposed the clusters from single-nucleus vagal sequencing onto the lung spatial analysis, we identified that all jugular and nodose clusters were spatially distributed in the airways and specific alveolar regions (**Figure 4C-E**). Specifically, the jugular and nodose clusters were predominantly found in areas surrounding the airways, where they overlayed with ciliated/club cells (**Figure 4E-H**). The nodose cluster were highly prevalent in alveolar regions, overlapping with alveolar cell types, including mesenchymal alveolar niche cells (MANC) (**Figure 4E-H**). The innervated cell types by the vagal clusters were identified as club cells and neuroendocrine cells. This is consistent with previous data characterizing these cells with immunohistochemical analyses(22, 43).

## Discussion

Vagal ganglia are key multimodal sensors of the lungs. As a multitude of the neuron types housed in the paired nodose and jugular ganglia innervate the lung, as demonstrated both physiologically and transcriptomically(3, 18, 44), their susceptibility to changes in response to asthma should be profound. Our study provides the first comprehensive transcriptomic analysis of vagal ganglia in response to asthmatic induction and correspondingly identifies cell types associated with neural patterns in the lung. Additionally, our data show the changes between control and asthmatic vagal ganglia. Furthermore, our single-nucleus and spatial mapping identify vagal clusters that innervate the lung and provide a map to identify the spatial locations of these clusters in the lung. We also show changes in specific vagal clusters between asthma and control congruent with bulk RNA-seq. Our results allow for a number of functional predictions to be made, but importantly, also provide tools for direct experimental strategies to address their morphology, physiology, connectivity, and function in response to allergic sensitization.

Our transcriptomic analysis of lungs and vagal ganglia, in tandem, further underscores the importance of vagal surveillance of the lungs. Bulk sequencing of the lungs revealed GO and KEGG pathway upregulation in *A. alternata* lungs compared to PBS controls which are typically associated with a neural phenotype. Of note, *Mrgprg* (6.99 Log2FC), *Runx1* (1.06 Log2FC), *P2RX2* (1.99 Log2FC), *Cacna1* (2.48 Log2FC), *Ntrk1* (2.00 Log2FC) were highly and significantly upregulated in the lungs in response to *A. alternata* induction. That nerve-associated receptors, ion channels, and nerve injury markers were increased in *A. alternata* lungs shows that vagal hypersensitization is a significant component of the asthmatic genotype/phenotype. This is consistent with an increase in sensory neuron hypersensitization and likely their influence in innervating neuroendocrine cells and the epithelium. In particular, *Mrgprg* and associated receptors from the mas-related gene protein coupled receptor family are known to be involved in neural hypersensitization as well as immune cell activation(15, 45). That these receptor families were upregulated in the vagal and lung transcriptome underscores their importance in asthmatic hypersensitization and further underscores the importance of neural activation in asthma. In addition to this novel finding, the lungs also upregulate *Tgfb* which is consistent with an allergic phenotype. Genes that were downregulated also show how allergic sensitization affects the lung transcriptome, as GO:0031012 extracellular matrix are downregulated in the lungs of asthmatic mice, demonstrating a disruption of the epithelial barrier.

The upregulation of particular genes in the vagal ganglia as a result of *A. alternata* induction supports the importance of the neuroimmune sensing capabilities of the vagal ganglia. To establish these data, single nucleus RNA sequencing confirmed bulk sequencing findings and show how the clustering of specific genes occurs. Single nucleus sequencing resulted in 7 unbiased clusters which were consistent with previous analyses by others(18). Interestingly, when jugular and nodose clusters are combined, *A. alternata* increases gene expression in jugular and nodose clusters which is consistent with the predictions from Kupari et al. to be lung-innervating ganglia(18). The agreement between bulk and single-nucleus sequencing and differentially expressed genes between *A. alternata* and control mouse ganglia demonstrate the upregulation of immune features and specifically immune features associated with immune memory. The single nucleus sequencing is consistent with spatial transcriptomics and shows the rostral-caudal distribution of these clusters consistent with a nodose/jugular distribution(22).

The transcriptomic distribution within the lungs showed the diversity of vagal cell clusters within the lungs. The vagal cell clusters are consistent with the physiologic discernment of lung cells and the single-cell lung atlases publicly available(46, 47). Interestingly, when we superimpose the information from single-nucleus sequencing of the vagal ganglia onto the lung spatial transcriptome map, we see the distinct distribution of these clusters within a finite space. The vagal cell clusters are distributed within the airways and distinct alveolar regions and were seen to ‘innervate’ club cells/neuroendocrine-like cells. To our knowledge, this is one of the first transcriptomic analysis to corroborate the functional descriptions from previous investigations. The most robust changes in response to *A. alternata* induction occurred in both jugular and nodose cells. This is consistent with the upregulation of gene types in the single nucleus and spatial vagal ganglia sequencing and the majority of changes occurring with immune transcripts.

Based on the cross-activation of genesets within the vagal ganglia and lungs and the close spatial profiles of vagal cluster sets within the lungs, our data show how sensory neurons are transcriptomically affected by allergic sensitization. As our data show consistency to previously published gene sets in the vagus, the changes seen in response to asthmatic induction are confined to certain cell types that are specific to lung-innervating cell types(3, 18). Our data are among the first to corroborate the functional purpose of these cell clusters and give a genetic basis to potential sensitization in response to asthmatic induction. Our data provide a rich dataset to test the functional consequences of allergic sensitization of vagal cell subsets.

## Methods

### Mice

All animal experiments were approved by The Lundquist Institute at Harbor UCLA Institutional Animal Care and Use Committee protocol # 32183. Mice were housed in a specific-pathogen-free animal facility at The Lundquist Institute. C57BL/6J mice were purchased from Jackson Laboratories. At the start of experiments, the mice were 5-8-weeks old. Age-matched male and female mice were used for experiments. In accordance with our animal protocol, all mice were anesthetized with 5% isoflurane and exsanguinated, followed by immediate dissection of tissues.

### *Alternaria alternata* asthmatic induction

*A. alternata* extract was purchased from CiteQ Laboratories (09.01.26) and dissolved to yield a final concentration of 25µg/ml in PBS. Mice were inoculated intranasally with 50µl on days 0, 3, 6, 9 where experiments took place 16 hours later(48). All control mice received PBS in the same volume intranasally.

### Immunoglobulin analysis

Immunoglobulin E (IgE) levels in serum were measured with enzyme-linked immunosorbent assay (ELISA, Thermofisher 501128838) according to the manufacturer’s instructions and analyzed with a BioTek Synergy H1 plate analyzer. Samples were compared using two-sided unpaired t-test.

### Histology

Lungs or vagal ganglia were embedded in paraffin and cut on a microtome (Leica) 5µm and stained with hematoxylin (Epredia 6765001) and eosin (Epredia 6766007or periodic acid, schiffs reagent (Sigma395B-1kit). Histological slides were mounted with permount toluene solution. Stained area was assessed with Image J, cell counts were normalized to length of tissue. Samples were compared using two-sided unpaired t-test.

### Bulk RNA sequencing

Vagal (nodose/jugular) ganglia were dissected from *A. alternata* and PBS mice. Ten vagal ganglia from five mice were pooled together per sample. A total of n=3 samples per group, per sex were obtained. Three sets of lungs (n=3) were used per group. RNA was isolated using the inTron Easy spin Total RNA extraction Kit (Boca Scientific 17221). RNA purity was assessed as >2.00, 260/280 ratio with spectrophotometry. RNA integrity was assessed prior to sequencing using the Agilent bioanalyzer and samples with RIN >7 were used for library construction. RNA sequencing was performed by Novogene Corporation Inc. (Sacramento, USA). mRNA was purified from total RNA using poly-T oligo-attached magnetic beads. To generate the cDNA library, the first cDNA strand was synthesized using a random hexamer primer and M-MuLV Reverse Transcriptase (RNase H−). Second-strand cDNA synthesis was subsequently performed using DNA Polymerase I and RNase H. Double-stranded cDNA was purified using AMPure XP beads and the remaining overhangs of the purified double-stranded cDNA were converted into blunt ends via exonuclease/polymerase. After 3’ end adenylation, a NEBNext Adaptor with a hairpin loop structure was ligated to prepare for hybridization. To select cDNA fragments of 150–200 bp in length, the library fragments were purified with the AMPure XP system (Beckman Coulter, Beverly, USA). Finally, PCR amplification was performed, and PCR products were purified using AMPure XP beads. The samples were sequenced on an Illumina NovaSeq 6000 with ≥20 million read pairs per sample. A full RNA-seq analysis pipeline was conducted by The Novogene Corporation which included alignment with HISAT2(49), PCA on the gene expression values (FPKM), differential gene expression (DEG) analysis using DESeq2(50), and finally a functional analysis including Gene ontology (GO) and Kyoto Encyclopedia of Genes and Genomes ontology (KEGG).

### Single-nucleus RNA sequencing

Vagal ganglia were dissected from *A. alternata* and PBS mice. Ganglia were immediately snap frozen in -80c. Eight-to-ten ganglia (from 4-5 mice) were pooled together per group; n=2 samples per group. Nuclei were isolated with the Minute™ single nucleus isolation kit (Invent biotechnologies inc #BN-020). The isolated nuclei were purified, centrifuged, resuspended in PBS with RNase Inhibitor, and diluted to 700 nuclei/µl for standardized 10x capture and library preparation protocol using 10x Genomics Chromium Next GEM 3’ Single Cell Reagent kits v3.1 (10x Genomics, Pleasanton, CA). Libraries were sequenced using an Illumina NovaSeq 6000 (Illumina, San Diego, CA, USA). The libraries were sequenced with ∼150 million PE50 reads per sample on Illumina NovaSeq. The raw sequencing reads were analyzed with the mouse reference genome (mm10) using Cell Ranger v7.1.0. To further clean the data, Cellbender was used to remove background noise(51). We employed a rudimentary cell quality control after CellBender, that is, removing cells using percentile-based thresholds on UMI count, gene count, and mitochondrial read fraction. UMAPs were created after (1) finding highly variable genes using the seurat_v3 algorithm, (2) normalizing counts per cell, (3) log scaling counts, (4) scaling counts of 2,000 highly variable genes and (5) performing PC analysis on those scaled values for the highly variable genes. A nearest-neighbor graph was constructed with 20 neighbors based on cosine distance in PC space (top 25 PCs).

### Spatial RNA sequencing

Lungs were dissected, flushed with PBS and perfused with 4% paraformaldehyde with 20cmH20 pressure for 10 minutes, then further fixed at 4°C for 24hours. Vagal ganglia were fixed in 4% paraformaldehyde for 2 hours. Tissues were processed and 5um paraffin slices were situated onto the xenium slide. DNA probes from a custom-selected DNA panel (**Supplementary data 4**), informed by bulk RNA-seq and single-nucleus RNA-seq. Gene sets were hybridized by ligation hybridization and amplification then subsequent rolling circle amplification. This process provides a robust spatial resolution and is based on MERFISH spatial sequencing therefore reducing false positives as ligation will not occur without complete probe binding. Subsequent series of hybridization, imaging, and probe removal were conducted to complete all probes. Downstream data visualization occurred via the Xenium imaging analyzer. Further analysis of spatial sequencing was completed with Seurat and Xenium Ranger v1.7.1.

### Bioinformatic analysis

The R-package Seurat (version 5.0.3) was used for the majority of the single-nuclei data analysis. The four independent biological replicates generated a total of 7670 sequenced cells with an average of approximately 117,958 reads, 613 genes and 910 UMIs detected per nuclei. All genes expressed in less than three nuclei were removed and the filtered gene barcode matrices from the individual runs were merged together. The cell-gene matrix was further filtered for mitochondrial-DNA derived gene-expression (20% was set as the high cut-off), the number of genes detected per cell (> 200 as low and < 5000 as the high cut-off), and the number of UMIs detected per gene (> 400 as the low and <10000 as the high cut-off) removing 1,322 nuclei. Counts were normalized using the SCTransform function, which applies a regularized negative binomial regression model to account for technical noise and nuclei-specific biases. This normalization step adjusts for differences in sequencing depth and captures biological variability across cells. Normalized data were used for principal component analysis (PCA) to reduce the dimensionality of the dataset. Uniform Manifold Approximation and Projection (UMAP) and t-distributed stochastic neighbor embedding (TSNE) were applied to visualize the cells in a low-dimensional space, capturing similarities between cells based on their gene expression profiles. The clustering of nuclei was performed using the original Louvain algorithm on the PCA-reduced data, defining clusters based on shared transcriptional profiles. Differential expression analysis was conducted to identify genes that were differentially expressed between nuclei clusters using the standard Wilcoxon rank sum test. Genes showing significant differential expression (adjusted p-value < 0.05) and high fold change (logFC > 0.25 or <-0.25) were considered as cluster-specific markers and used to characterize distinct cellular populations based on their biological functions and expression profiles. Clusters were annotated based on known marker genes and enriched biological functions using gene ontology analysis. Cell type identities were assigned by comparing marker genes with reference datasets(18). Clusters and gene expression patterns were visualized using feature plots, violin plots and dot plots to illustrate differential expression and cluster-specific marker genes.

### Correlation analysis of bulk and single nuclei

We utilized Transcript per Million (TPM) values from bulk RNA sequencing data and Counts Per Million (CPM) values from single-cell RNA sequencing data for our analysis. To assess the relationship between these datasets, we calculated the correlation between TPM values from the bulk RNA sequencing and Cluster PM values from the single-nuclei RNA sequencing. This approach allowed us to evaluate the consistency and coherence of gene expression profiles across different types of RNA sequencing data. To evaluate the correlation between our dataset and the previously published Kupari dataset, we first calculated the CPM for each dataset separately, followed by log2 transformation of the CPM values. We then performed a correlation analysis on the log2-transformed CPM values.

### Cellchat

CellChatv2.1.1 was used to elucidate the intercellular communication networks in our single-nuclei RNA-seq dataset(52). We employed the full CellChatDB, excluding the ‘Non-protein Signaling’ category, for our cell-cell communication analysis. This tool utilizes known ligand-receptor interactions to infer communication patterns between different cell types. CellChat objects were created from our Seurat objects using the createCellChat function. We then followed the standard CellChat pipeline, which involved computing the communication probability using the computeCommunProb and computeCommunProbPathway functions. Subsequently, we calculated network centrality scores and the contribution of known ligand-receptor pairs using the netAnalysis_computeCentrality and netAnalysis_contribution functions, respectively. The cell communication results were visualized using the standard functions provided by the CellChat package.

### Deconvolution of Xenium lung and neuron dataset

To deconvolute our Xenium data from the lung and ganglia, we utilized the RCTD (Robust Cell Type Deconvolution) method inside of the spacexr package (v2.2.1) in conjunction with appropriate reference datasets(53). For the lung Xenium data, we employed a dataset on LungMap(54, 55), which provided a detailed reference for lung cell type-specific expression profiles. We integrated the LungMap dataset with the lung Xenium data using the create.RCTD function with the LungMap dataset as the reference and our lung Xenium data as the query. RCTD’s deconvolution algorithm was then applied using the run.RCTD function to estimate the proportion of each lung cell type in the Xenium samples, allowing us to accurately infer cell type compositions. Subsequently, the ImageDimPlot function was used with the group.by parameter set to “predicted.celltype” to visualize the predicted cell types overlayed on our lung Xenium section.

For the vagal ganglia Xenium data, we integrated our single-nuclei RNA sequencing dataset with the published Kupari et al.(18) dataset. This integrated dataset served as a reference for cell type-specific expression profiles relevant to ganglia. Using this combined reference, we applied RCTD to the ganglia Xenium data to deconvolute the cell type proportions. The deconvolution process involved comparing the expression patterns observed in the Xenium data with the reference matrix to identify the cellular composition. To ensure the robustness of our findings, we performed cross-validation and sensitivity analyses to validate the stability and reliability of the deconvolution results for both tissue types. Additionally, we used our ganglia single-nuclei dataset integrated with the Kupari et al. dataset as a reference for the RCTD algorithm to see if we could identify ganglia cells innervating the lung.

## Data availability

Bulk, single-nucleus, and spatial RNAseq raw data can be accessed via PRJNA1169017. All data reported in this paper will be shared by the lead author upon request. Any additional information required to reanalyze the data reported in this paper is available from the lead author upon reasonable request.

## Acknowledgments

NJ is supported by NIH grant R21AI159221, R56AI175328, UCLA CTSI UL1TR001881-01, and T32KT4708 of the Regents of the University of California Tobacco-Related Diseases Research Program. T.A.D. is supported by NIH AI171795 and Veterans Affairs BLR&D BX005073. WY is supported by NIH grants P50HD098593 and R01HD099924. Figure 1a was completed with Biorender under a CC-BY-NC-ND license.

