## Supplementary figures for "Comprehensive sequencing of the lung neuroimmune landscape in response to asthmatic induction"

**Supplementary Figure 1.**

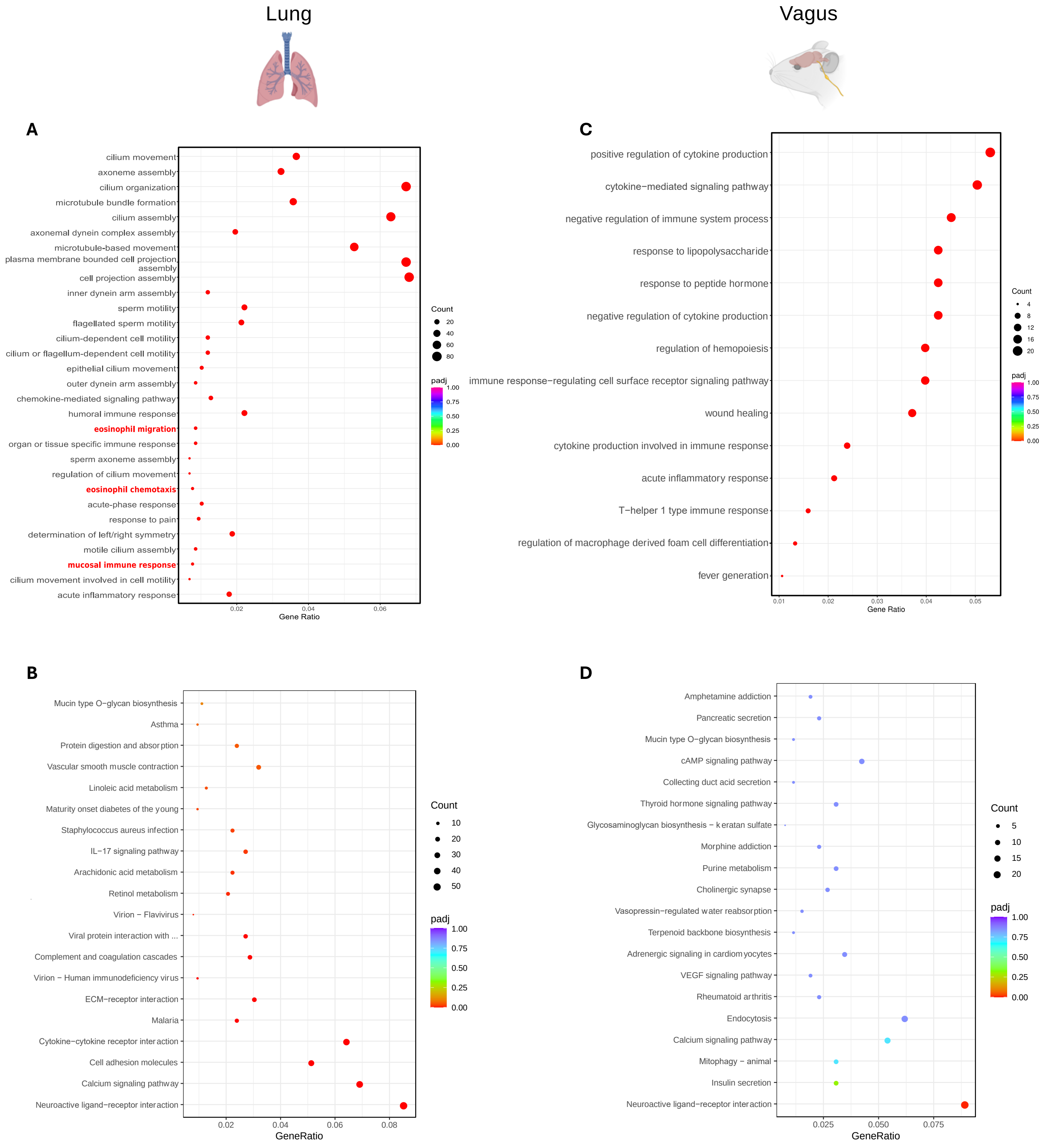


**Supplementary figure 1. GO and KEGG analysis. LUNG: A)** GO dot plot showing enriched processes differentiating *A. alternata*-induced asthma conditions from control. **B)** KEGG dot plot highlighting pathway enrichment differences between *A. alternata*-induced asthma from control. **C)** GO dot plot showing enriched processes differentiating *A. alternata*-induced asthma conditions from control. **D** KEGG dot plot highlighting pathway enrichment differences between *A. alternata*-induced asthma from control.

**Supplementary Figure 2.**

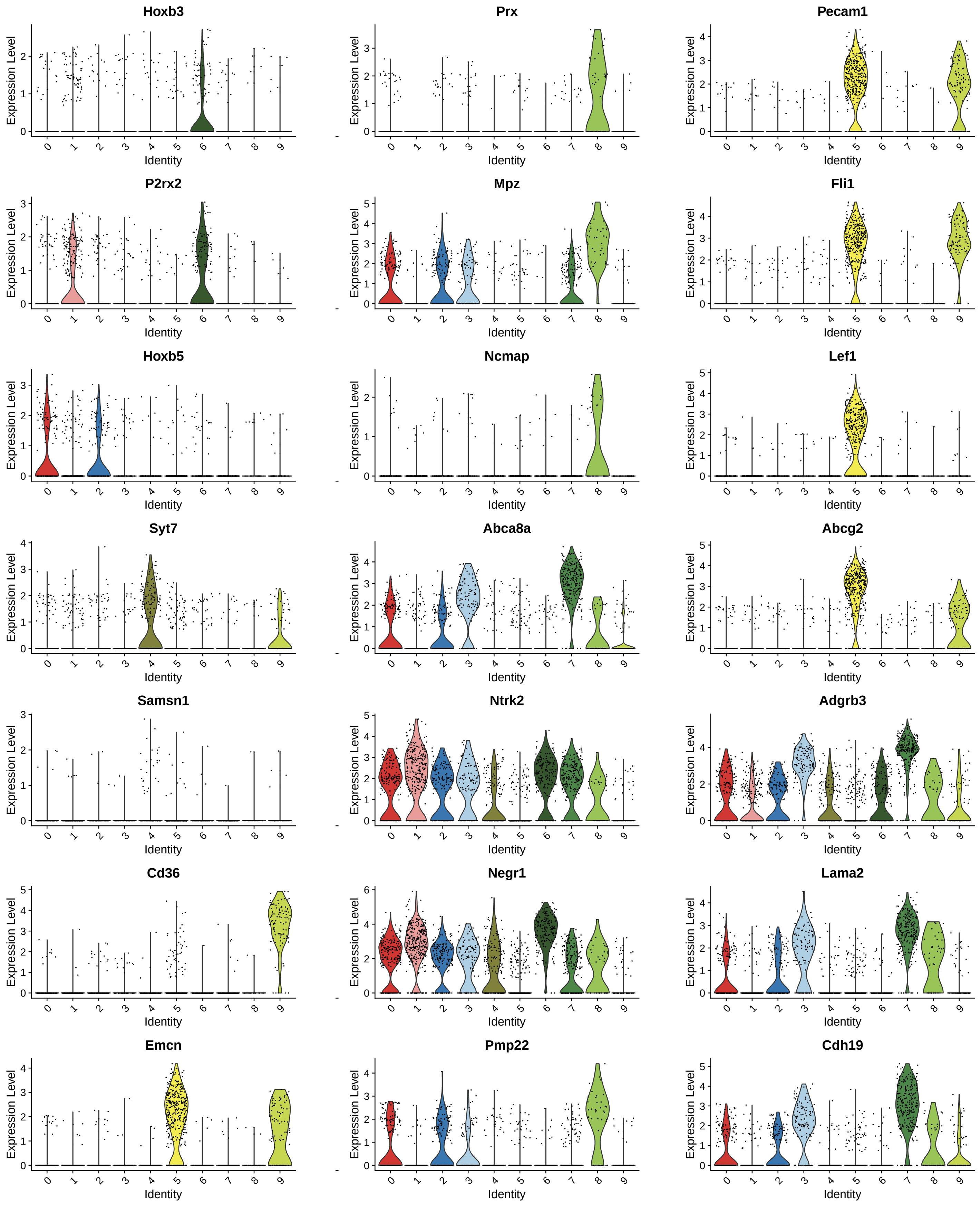


**Supplementary figure 2. Canonical gene markers identifying cell types in single-nuclei RNA-seq dataset a)** Violin plots showing the expression of key marker genes in the 11 unbiased clusters.

**Supplementary Figure 3.**

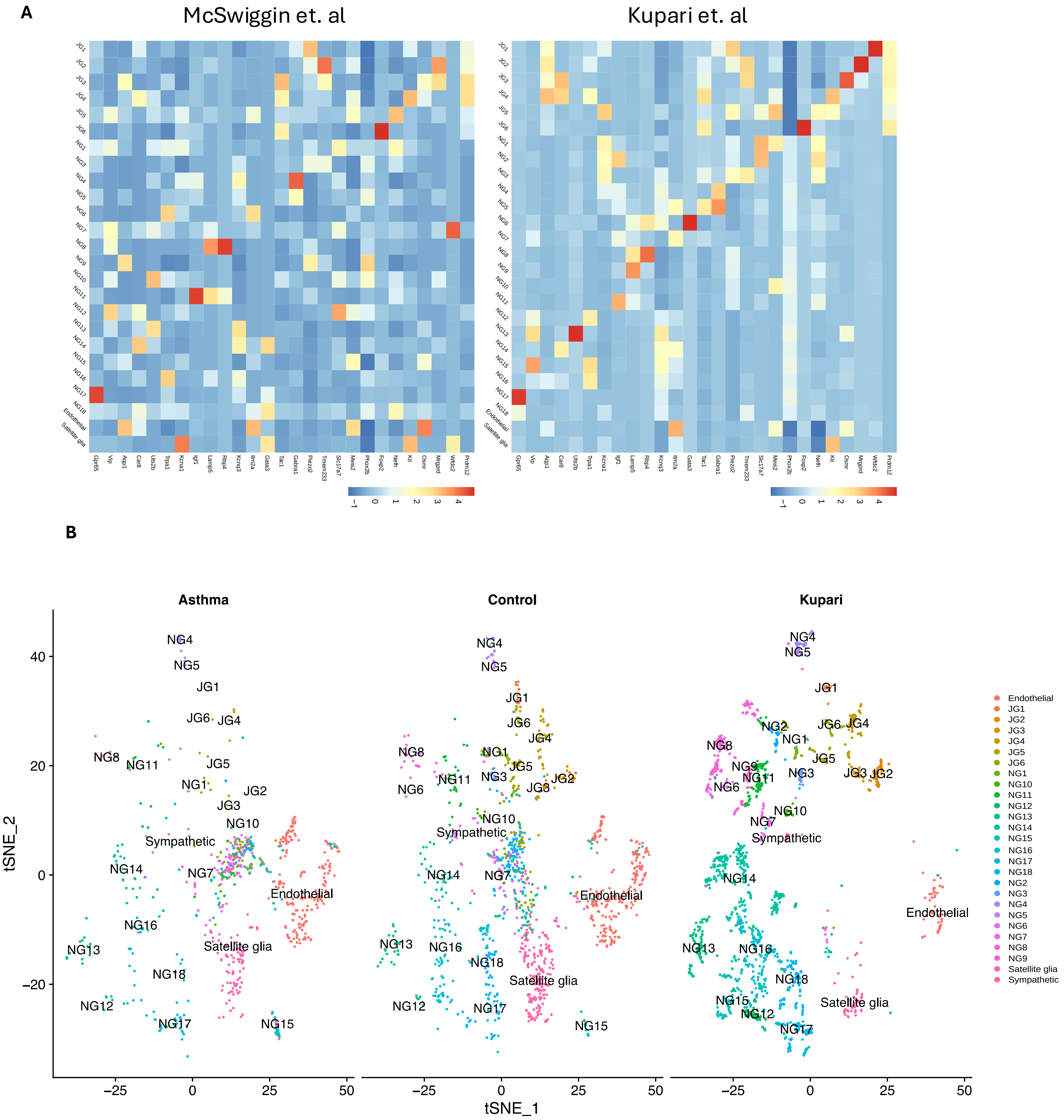


**Supplementary figure 3. Single-nucleus sequencing of control and asthmatic vagal ganglia are similar to previous single-cell sequencing of healthy vagal ganglia. A)** Heatmap of showing expression of key genes from the data provided by Kupari et al. in the current dataset **B**) Tsne reveals a high similarity between the current single-nucleus analysis to the single cell analysis of Kupari et al.

#### Supplementary Figure 4.

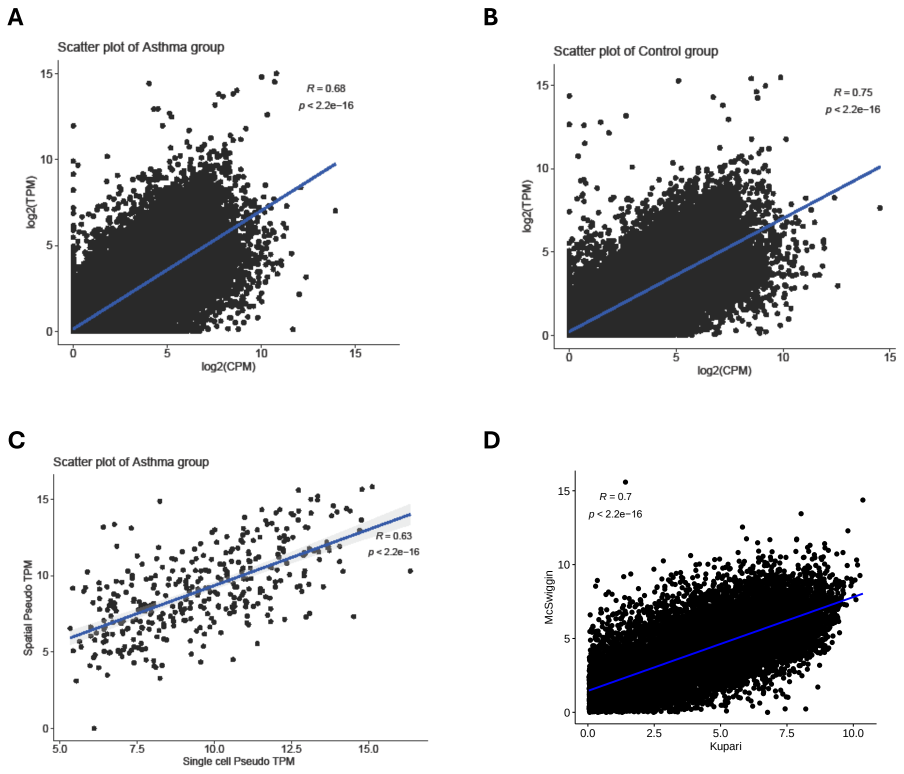


**Supplementary figure 4. Correlation between datasets. A)** Correlation between bulk and single nucleus sequencing of vagal transcripts in *A. alternata*. **B**) Correlation between bulk and single nucleus sequencing of vagal transcripts in control. **C**) Correlation between single nucleus sequencing and spatial sequencing of vagal transcripts. **D)** Correlation between the current single-nucleus dataset and previously published single-cell dataset from Kupari *et. al*.

### Supplementary Figure 5.

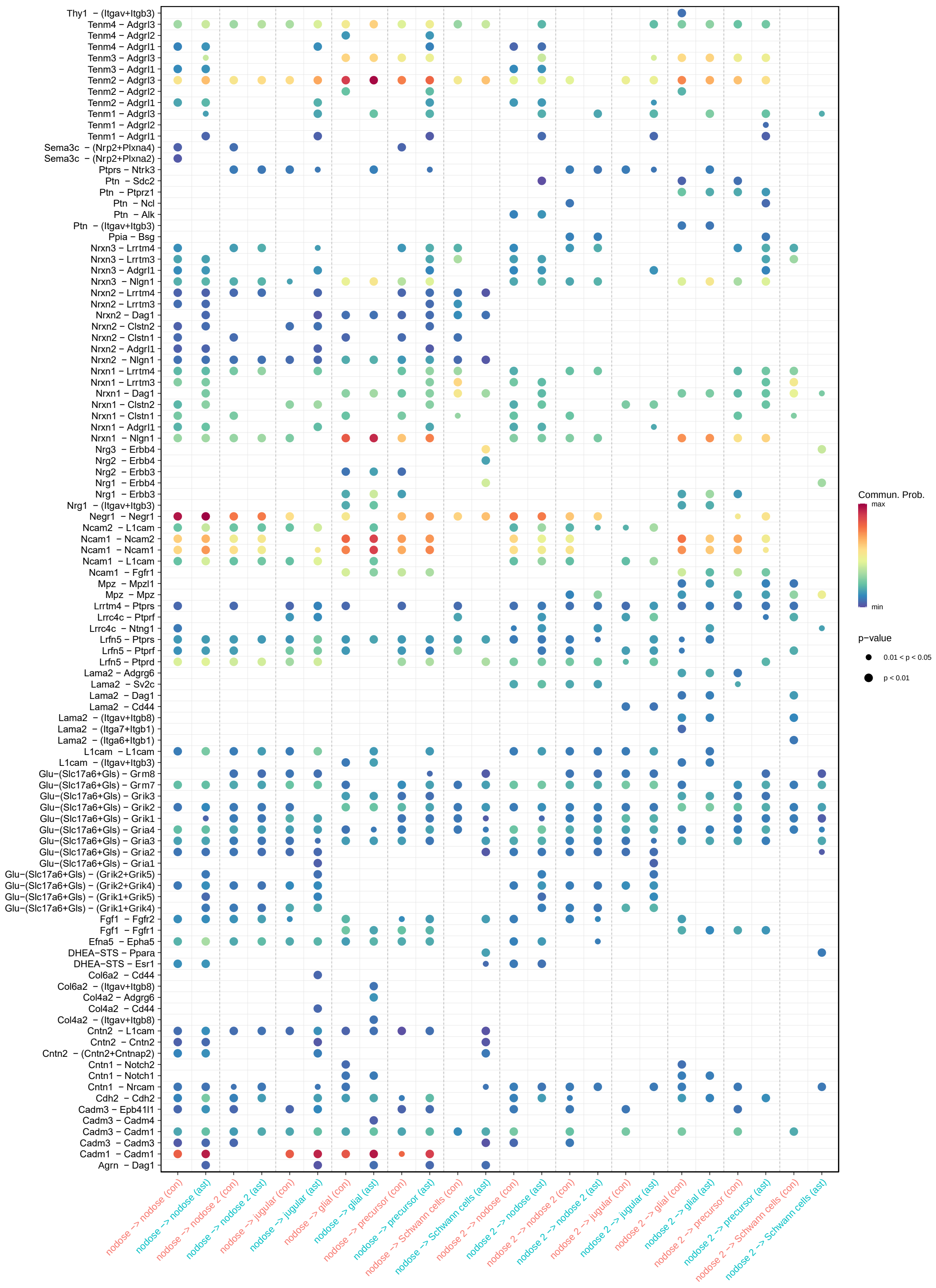


**Supplementary figure 5. Differential CellChat expression.** Differential expression of ligands and receptors between *A. alternata* and control vagal transcripts. Jugular was the source and all other cell types were targets.
